# SeqVerify: An accessible analysis tool for cell line genomic integrity, contamination, and gene editing outcomes

**DOI:** 10.1101/2023.09.27.559766

**Authors:** Merrick Pierson Smela, Valerio Pepe, George M. Church

**Affiliations:** Wyss Institute at Harvard University, Cambridge, Massachusetts, United States of America; Department of Genetics, Harvard Medical School, Harvard University, Cambridge, Massachusetts, United States of America

**Author notes:** Equal contributions (Merrick Pierson Smela, Valerio Pepe).

**Keywords:** Stem cell, Pluripotent stem cell, Whole genome sequencing, microbial contamination, aneuploidy, genome editing, single nucleotide polymorphisms, software, quality control, SeqVerify

## Abstract

Over the last decade, advances in genome editing and pluripotent stem cell (PSC) culture have let researchers generate edited PSC lines to study a wide variety of biological questions. However, abnormalities in cell lines such as aneuploidy, on-target and off-target editing errors, and microbial contamination can arise during PSC culture or due to undesired editing outcomes. Any of these abnormalities can invalidate experiments, so detecting them is crucial. The ongoing decline of next-generation sequencing prices has made whole genome sequencing (WGS) an effective quality control option, since WGS can detect any abnormality involving changes to DNA sequences or presence of unwanted sequences. However, this approach has suffered from a lack of easily usable data analysis software. Here, we present SeqVerify, a computational pipeline designed to take raw WGS data and a list of intended edits, and verify that the edits are present and that there are no abnormalities. We anticipate that SeqVerify will be a useful tool for researchers generating edited PSCs, and more broadly, for cell line quality control in general.

## 2 Introduction

Pluripotent stem cells (PSCs) have found important uses in many areas of biological research, and their ability to differentiate into a variety of cell types has enabled the development of cell-based therapies derived from PSCs. Gene editing technologies such as CRISPR-Cas9 have enabled the engineering of PSC lines containing specific alleles of interest, such as disease-relevant mutations or fluorescent reporters.

However, over the years, researchers have identified several common abnormalities that can arise during PSC culture. First, PSCs can become aneuploid due to chromosomal rearrangements or mis-segregation. The most frequent aneuploidies in PSC cultures involve chromosomal or sub-chromosomal duplications^1,2^, and some of these, such as gain of 20q11.21, have been characterized to affect the phenotypes of the cells^3,4^. Such aneuploidies are relatively common, affecting roughly twelve percent of tested PSC lines on average^2^, with the frequency increasing over long-term passaging.

Second, PSCs can gain point mutations, which can be enriched during prolonged culture due to providing a growth advantage. For example, the tumor suppressor genes *TP53* and *BCOR* are recurrently mutated in PSCs^5,6^. Although mutation rates in PSCs are not abnormally high^7,8^, harmful mutations can often be present in somatic cells used to derive induced PSCs^6^, or can occasionally arise during PSC culture^5^.

Third, as with other cell cultures, PSC cultures can be contaminated with microbes such as *Mycoplasma*. This is a relatively common problem; a study in 2015 was able to detect sequencing reads mapping to *Mycoplasma* in eleven percent of mammalian cell culture datasets in the NCBI Sequence Read Archive^9^. Good cell culture practice, including recurrent testing, is essential for avoiding contamination.

Furthermore, gene editing of PSCs can introduce additional abnormalities. At the on-target site, undesired editing outcomes may be present. Traditional PCR-based genotyping can sometimes fail to detect these outcomes when they involve a large insertion of plasmid or mitochondrial DNA into the target site^10^. Although off-target editing is usually rare^11^, the rate may be greatly increased when less specific editing tools, such as APOBEC-based cytosine base editors, are used^12,13^.

In order to ensure valid experimental results, and the safety of PSC-derived therapeutics, it is important to detect these abnormalities and choose PSC lines without them. Existing quality control methods, including karyotyping, SNP arrays, and quantitative PCR, typically focus on detecting one particular type of abnormality ^1^. However, the ongoing decline of next-generation sequencing prices has made whole genome sequencing (WGS) an effective quality control option. Notably, WGS is an all-in-one detection method for any abnormality involving changes to DNA sequences (such as aneuploidy or mutations), presence of unwanted sequences (such as plasmid integration or *Mycoplasma*), or cell line misidentification. WGS data can also help select PSC lines for experiments based on polygenic risk scores for traits of interest ^8^

Yet until now, WGS analysis has required considerable expertise in bioinformatics due to a lack of easily usable software. Although other researchers have used WGS for PSC quality control,^8^ this only looked at wild-type cell lines and the analysis pipeline code was not publicly available. Here, we present a computational pipeline, SeqVerify, that analyzes short-read WGS data for quality control of wild-type or edited PSCs. SeqVerify can validate on-target genome editing, find the insertion sites of untargeted transgene integrations, and detect mutations, aneuploidies, microbial contamination, and misidentification. SeqVerify provides an end-to-end analysis framework, with simple inputs (raw WGS data and a list of intended edits) and easily interpretable outputs. We have made our pipeline easily installable via Bioconda. Furthermore, we validate the performance of SeqVerify on a set of knock-in hiPSC lines generated in our lab. We anticipate that WGS and SeqVerify will be a valuable quality control method for researchers working with PSCs, and more broadly, for cell line quality control in general.

## 3 Results

### 3.1 The SeqVerify Pipeline

#### 3.1.1 Overview of the Seqverify Pipeline

SeqVerify is an end-to-end pipeline that performs a variety of quality control functions (Figure 1). First, SeqVerify generates an “augmented genome” from a reference genome and a user-provided list of targeted edits and/or untargeted transgene insertions. SeqVerify will then align the raw WGS reads to this augmented genome, validate edits, detect insertion sites, and analyze copy number variation. SeqVerify also detects microbial contamination using KRAKEN2^14^. Additionally, SeqVerify will align the reads to a wild-type reference genome, detect and filter single-nucleotide variants (SNVs), and annotate them using the ClinVar database^15^. Finally, if two or more samples are analyzed using SeqVerify, the SNVs can be automatically compared. This is useful for detecting cell line misidentification, or for identifying SNVs arising during cell culture or editing which were not present in the original cells.

**Figure 1:**
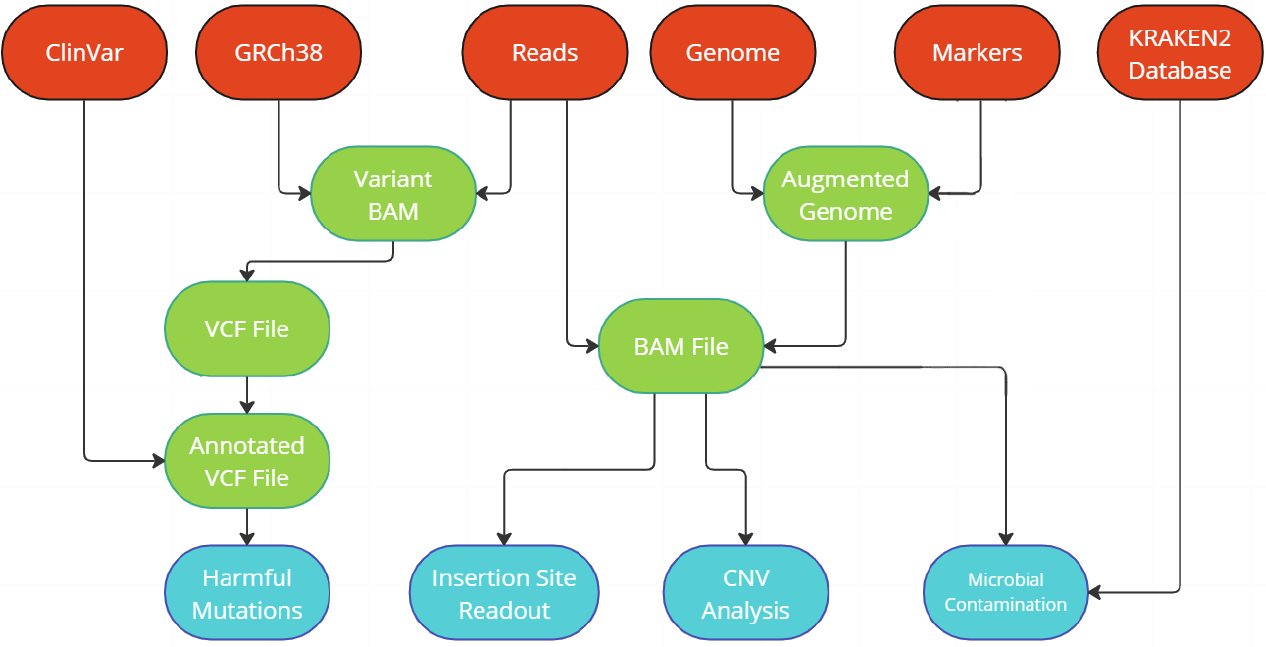
A Mlowchart showing the steps involved in the SeqVerify pipeline. In red, the possible types of inputs that the pipeline can take for the types of analysis that can be performed. In light green, the major intermediate files produced during the running of the pipeline, output in the non-temporary output folder. In light blue, the four major outputs of the pipeline: the insertion site readout and CNV analysis graphs and binaries, as well as the microbial contamination and harmful mutation readouts if the KRAKEN and SNV Analysis portions of the pipeline are enabled, respectively.

#### 3.1.2 Installation

SeqVerify was developed for and tested on Linux systems (including Windows Subsystem for Linux). It can also run on MacOS or other Unix-based systems, although automatic IGV plots do not currently work on MacOS. SeqVerify can be installed using the conda package manager, and it can be downloaded from Bioconda with the following command:

~~~
conda install -c bioconda seqverify
~~~

We recommend this installation method since it also comes pre-packaged with all the dependencies needed to run. However, it can also be downloaded from GitHub with dependencies installed separately.

In terms of technical specifications, any system powerful enough to run BWA-MEM in reasonable time will also be acceptable for SeqVerify, and there are options available for multithreading and limiting memory usage that permit users to tune SeqVerify to their needs. The overall runtime using 20 threads on an Intel® Xeon® Processor E5-2683 v4 (40M Cache, 2.10 GHz) is approximately 11-12 hours per sample at 10X genome-wide coverage, increasing proportionally with increasing sequencing depth. Detailed usage instructions for SeqVerify are provided in Supplementary File 1.

#### 3.1.3 Automatic Download of Reference Data

The `seqverify --download_defaults` command will automatically download all the default files for a standard analysis of human cells. These are:

- T2T-CHM13v2.0 as the overall reference genome,^16,17^
- GRCh38 as the reference genome for Single Nucleotide Variant calling,
- PLUSPF 8GB as the default KRAKEN2 database,
- ClinVar as the default VCF annotation database,
- snpEff.config as a fresh snpEff configuration file should the user want to manually specify advanced snpEff options.

SeqVerify downloads all of them from their respective FTP servers at once and stores them in a `seqverify_defaults` folder in the working directory where the command is run. If, when running the pipeline, certain options are left blank (reference genome, KRAKEN database, etc.), SeqVerify will automatically attempt to use these from the `seqverify defaults` folder to correctly run the pipeline.

#### 3.1.4 Validation of Edits at Known Target Sites

To validate edits at known sites, SeqVerify takes an input file listing genomic coordinates and DNA sequences to be inserted, edited, or deleted at those coordinates. SeqVerify will use this information to generate an edited reference genome, corresponding to the user’s intended edits. After aligning reads to this genome using BWA-MEM^18^, SeqVerify will automatically generate figures using IGV^19^, displaying the genomic coordinates provided by the user and showing the aligned reads. Since SeqVerify saves the BAM file output after alignment, the user can also manually open this file in IGV if more detailed inspection is desired. We tested SeqVerify on hiPSC lines that we edited with fluorescent protein reporter knockins at loci such as *NANOS3* and *REC8*. On-target homozygous edits (Figure 2A) are easily distinguishable from undesired outcomes (Figure 2B).

**Figure 2:**
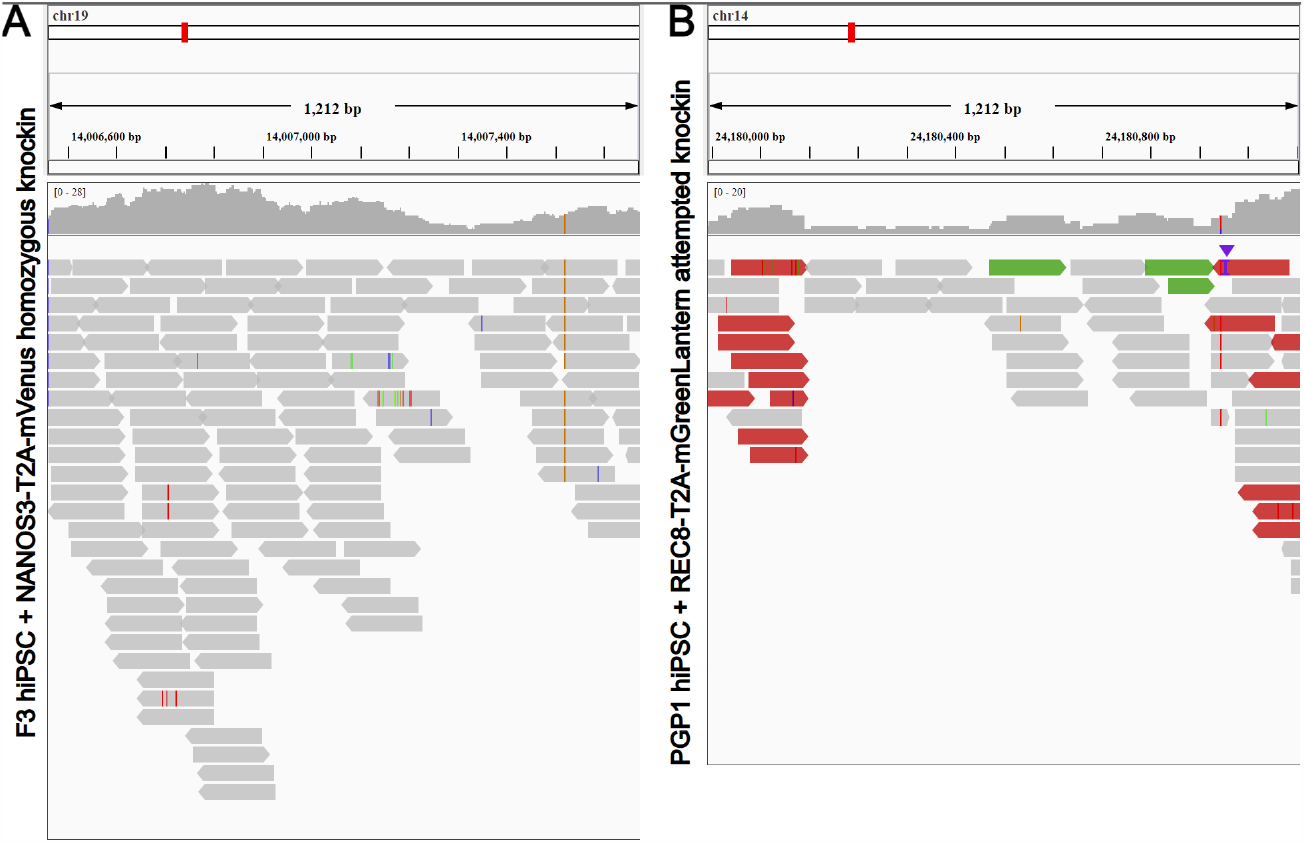
On-target edit validation using SeqVerify. hiPSCs with fluorescent reporter knockins (panel A: NANOS3-T2A-mVenus; panel B: REC8-T2AmGreenLantern) were analyzed by WGS and the SeqVerify pipeline. IGV plots were automatically generated showing the insertion and 200bp of Nlanking wildtype sequence on either side. The NANOS3 reporter line is a homozygous knockin. A SNP present in the starting cells is visible (brown line). By contrast, the REC8 reporter line is heterozygous; read pairs highlighted in red by IGV denote a ”deletion” of the T2A-mGreenLantern sequence on one of the alleles. Additionally, read pairs highlighted in green by IGV map both to the inserted sequence, and to bacterial plasmid DNA. This clearly shows an undesired editing outcome.

#### 3.1.5 Detection of Untargeted Transgene Insertions

SeqVerify will also accept untargeted transgene sequences as input. This is useful for finding the insertions of transposons and lentiviruses, or for detecting inadvertent integration of plasmids used in editing or iPSC reprogramming. User-provided sequences are appended to the reference genome and treated as extra chromosomes during alignment. Similarly to targeted insertions, SeqVerify will display reads aligning to these transgenes using IGV. We tested SeqVerify for the ability to detect undesired integration of our gene editing plasmid backbone in edited cells, an abnormality which is common yet easily missed by standard PCR genotyping^10^. In one of our cell lines edited at the *DDX4* locus, we observed plasmid integration into the target site (Figure 3A).

**Figure 3:**
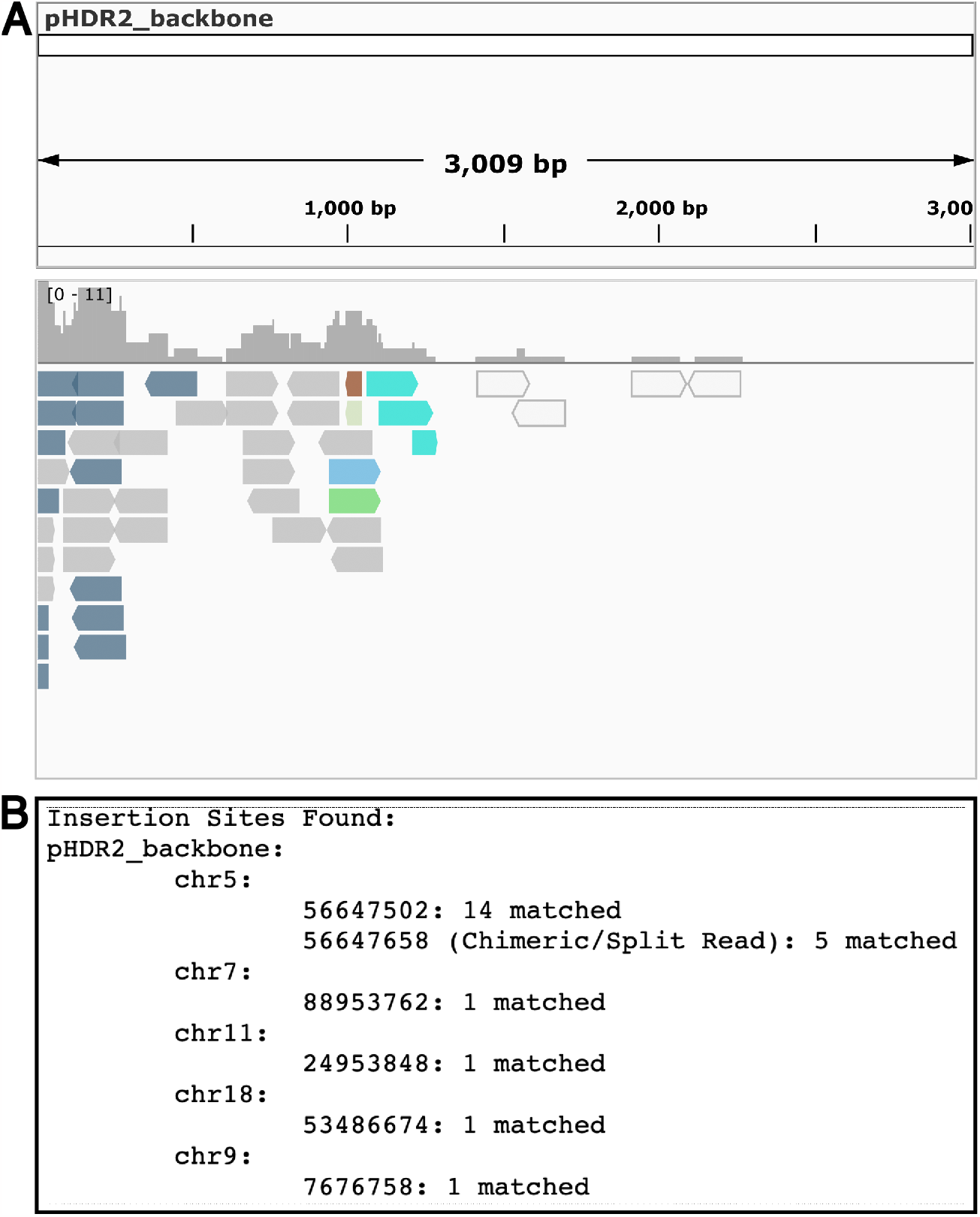
Detection of an undesired plasmid insertion in hiPSCs edited with a *DDX4* knockin. (A) The automatically generated IGV plot shows reads aligning to the plasmid backbone. Reads highlighted in blue have their mates mapped to the human genome. (B) Automatic detection of the insertion site (at the *DDX4* locus on chr5).

However, manually looking through alignments to detect insertion sites is tedious and does not scale well. Therefore, SeqVerify will also automatically find insertion sites by detecting junctions between transgenes and the host genome. After reads have been aligned to the modified reference genome, SeqVerify will parse the output SAM file, extract chimeric read pairs aligning to a transgene and to a host chromosome, and output a human-readable text file listing any detected insertion sites (Figure 3B).

#### 3.1.6 Copy Number Variation

For copy number variation (CNV) detection and analysis, SeqVerify uses the CNVPytor package^20^ and generates a Manhattan plot of the normalized read depth at a default resolution of 100kbp. These plots are useful for visualizing aneuploidies, as shown in Figure 4. We tested this by comparing WGS data from euploid hiPSCs (Figure 4A) and aneuploid HEK293 cells (Figure 4C). We also serendipitously detected a 12p duplication in one of our hiPSC samples (Figure 4B) that has been previously reported as a common abnormality in hPSCs^21^. Besides generating plots, CNVpytor will also call CNVs and output a list of any CNVs found in the sample.

**Figure 4:**
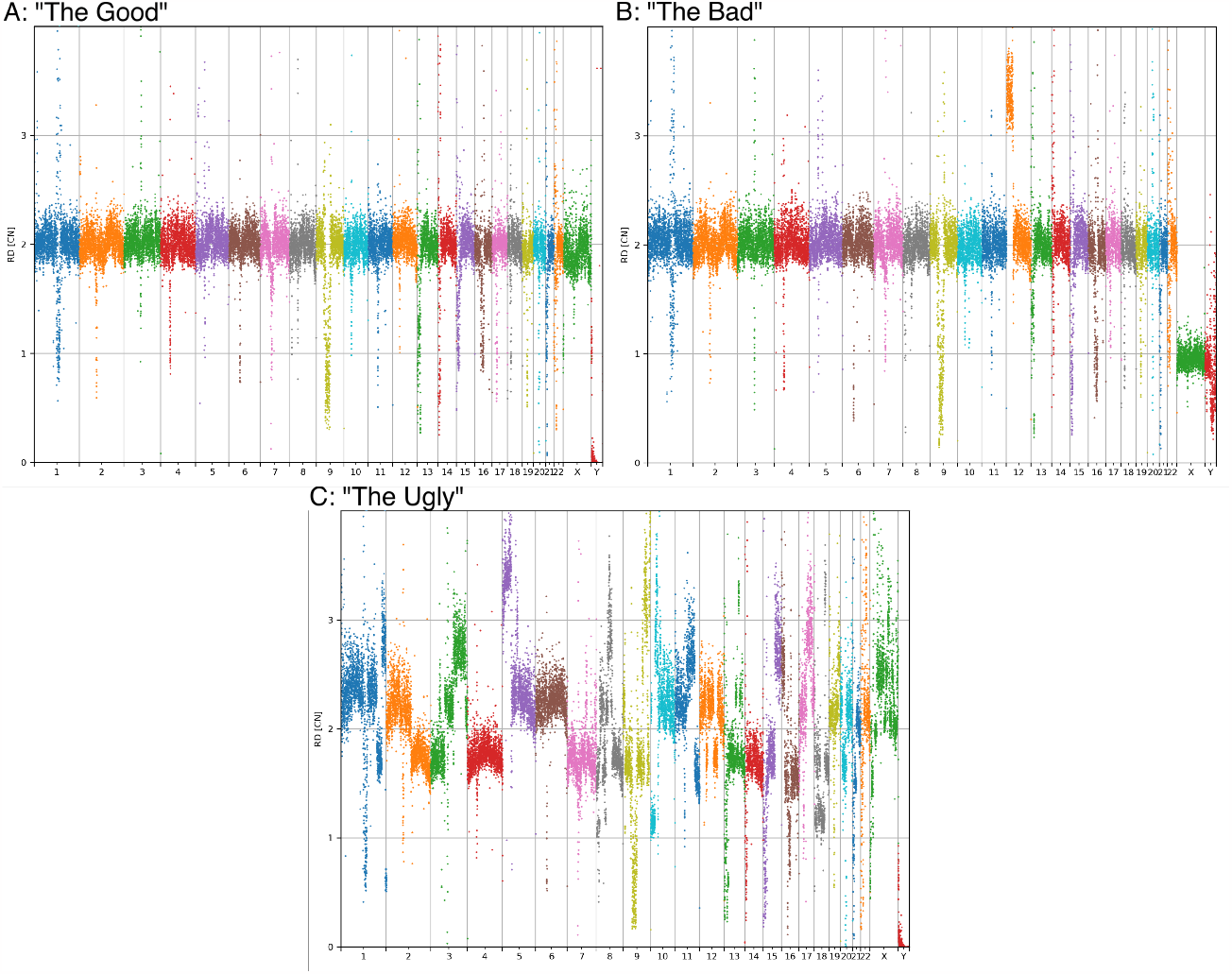
Aneuploidy detection using CNVpytor. WGS data from two hiPSC lines (panel A: female 46XX; panel B: male 46XY dup(12p)) and HEK293 cells (panel C) were analyzed using SeqVerify. CNVpytor plots are shown. Panel A shows a normal female karyotype, although some read depth variations are present in repetitive DNA near centromeres and telomeres due to challenges in aligning these sequences. Panel B shows an aneuploid male karyotype. Panel C shows the massive aneuploidy of HEK293 cells.

Additionally, transgene copy numbers can be estimated using the normalized read depth. Transgene plots are produced using an automatically generated IGV batchfile, which takes screenshots of genomic coordinates corresponsing to transgene sequences. This is helpful for distinguishing full vs. partial transgene insertions. However, taking automatic screenshots requires the XVFB package which is not available on all systems (for example, MacOS). Therefore, we developed an alternative plotting system using the Python library matplotlib. The internal plotting tool uses the output from running `samtools depth` on the data to compare the amounts of reads per 30bp to an estimate of an expected value of reads per 30bp bin calculated from the average read depth. It then plots this data as a histogram of copy number against bin, and makes one of these plots per transgene.

#### 3.1.7 Contaminant Detection

SeqVerify performs contaminant detection using KRAKEN2^14^. After aligning reads to the human reference genome, SeqVerify extracts all read pairs where either the read or its mate are unmapped, and runs KRAKEN2 to classify them. We recommend that a KRAKEN database containing human sequences be used (for example, PlusPF-8 which is used as default), since unmapped reads may still contain some human sequences which may otherwise generate false positives. Further analysis to estimate contaminant species abundance is then performed with BRACKEN^22^.

We validated this contaminant detection by running it on data from twelve hiPSC lines which tested negative for *Mycoplasma* by PCR-based methods, and one which tested positive. We found no signs of contamination (less than 50 reads for any bacterial or viral species) in the negative samples, and the positive sample had 23,891 reads mapping to *Mycoplasmopsis arginini* (Supplementary File 2). Furthermore, we analyzed previously published WGS reads from HEK293 cells (SRA number: SRR18054575) to see if this method could detect adenovirus 5, which was used to transform those cells in their original derivation from fetal tissue. We successfully detected 497 reads mapping to Adenoviridae, of which 104 specifically matched human adenovirus 5. Additionally, we found that the HEK293 cells used in that dataset were contaminated with *Mesomycoplasma hyorhinis* (Supplementary File 2).

#### 3.1.8 Single Nucleotide Variant Analysis

Single Nucleotide Variant (SNV) analysis requires re-alignment of the reads to an un-edited reference genome. Most human SNV annotation databases (for example, ClinVar) use hg38 instead of CHM13 as their reference genome. Therefore, in the SNV portion of the pipeline, the reads are re-aligned to hg38, with BCFTOOLS subsequently used to generate a VCF file of the variants found in the reads^23^. SeqVerify then annotates the VCF file using SnpEff and SnpSift and (by default) the use of the ClinVar clinical database for further annotation and loss of function and effect prediction^24,25^. Finally, SeqVerify then takes the annotated VCF file and filters it, generating a human-or machine-readable readout of all mutations above a certain quality score and severity threshold, their effects, genes, any loss of function, and their homozygosity, among other data. An example of such a readout is provided in Supplementary File 3, which also contains other example output.

#### 3.1.9 Single Nucleotide Variant Comparison

After running the main SeqVerify pipeline on at least two samples, the SNV results can be compared using the `seqverify --similarity` command. This classifies the SNVs according to whether they are shared between samples or specific to one sample. This is implemented using the `bcftools isec` command, with further processing to compute the concordance between the samples. This comparison is useful to detect potential cell line misidentification, or to detect mutations in edited cell lines that were not present in the original cells.

## 4 Discussion

In recent years, WGS has emerged as a powerful tool for genetic research and clinical diagnostics. The continued decline in sequencing prices over time has made WGS an attractive method, and we anticipate this decline will continue in the future. Remarkably, WGS (at 10X coverage) is now cheaper than karyotyping, and cost-competitive with commercial microarray-based services such as KaryoStat. The power of WGS was illustrated by a recent study which analyzed 143 wild-type hESC lines for SNVs and copy number variations, generating a database of sequence-verified hESC lines^8^. However, that study was limited to looking at variation in wild-type cells, and also did not publish alignment and variant calling code. Overall, the lack of convenient data analysis methods has presented a barrier for routine use of WGS in cell line quality control.

Therefore, we developed SeqVerify, a pipeline to analyze WGS data to validate genetic edits and check for abnormalities, including aneuploidy, mutations, and microbial contamination. SeqVerify is easily installable using bioconda, and provides a start-to-finish pipeline, taking raw sequencing reads as input and outputting quality control data. We have been using SeqVerify internally in our lab for the past year, and are now publishing it to share with others.

### 4.1 Comparison of WGS with previous quality control methods

#### 4.1.1 Validating on-target editing

PCR and Sanger sequencing is cost-effective for initial screening, but struggles to detect certain unwanted editing outcomes such as large insertions of plasmid or mitochondrial DNA^10^. Additionally, some transgene delivery methods, including lentivirus and PiggyBac transposons, involve the random insertion of transgenes into the host genome. Detecting the insertion sites of these transgenes is important for quality control to ensure that essential host genes are not compromised. Furthermore, if plasmids are introduced into cells for gene editing, unwanted plasmid integration events may occur. Specialized PCR-based methods can efficiently map insertion sites of known DNA sequences^26^, although they may miss partial insertions where the primer binding site is not inserted. WGS can identify insertion sites in an unbiased manner, although if multiple different transgenes are inserted using similar transposons, short-read sequencing cannot always definitively assign the transgene present at each insertion site.

#### 4.1.2 Detecting deleterious mutations

When establishing a new cell line, it is important to rule out the presence of deleterious mutations. These mutations may be due to off-target editing, or may arise spontaneously. Targeted amplification and sequencing of mutation hotspots or predicted off-target edit sites can provide some information, but may miss important mutations. WGS is the only practical way to detect mutations across the entire genome.

#### 4.1.3 Detecting aneuploidy

The traditional method of detecting aneuploidy is by karyotyping, with G-banding or FISH. This is effective, but laborious, requiring the preparation and staining of metaphase spreads. Additionally, smaller abnormalities, such as 20q11.21 duplication which is common in PSCs, may be missed with standard karyotyping methods^3^.

In recent years, DNA microarrays (for example, Thermo Fisher KaryoStat+) have also been used to detect aneuploidy. These are more sensitive, and require only a DNA sample. However, the decrease in cost of WGS over the last few years has made WGS a cost-competitive alternative for this method, with the added benefit of achieving even higher resolution. One drawback of short-read WGS (and especially microarrays) relative to karyotyping is that they are less sensitive at detecting balanced chromosomal translocations. If it is important to detect these translocations (and it may not be, given that they are rare, and typically have mild effects^27^, then traditional karyotyping or long-read sequencing should be used.

#### 4.1.4 Detecting microbial contamination

Testing for microbial contamination, especially *Mycoplasma*, is a critical part of good cell culture practice. This can be done by multiple methods, including PCR and enzyme-based kits. We do not recommend WGS as a routine screening method for contamination, due to higher cost, longer turnaround time, and lower sensitivity than PCR. Nonetheless, WGS data can be analyzed to check for the presence of microbial contamination. In principle, any contaminant with a DNA genome can be detected. This is an advantage over PCR-based methods, which can only detect contaminants matching the PCR primer pairs used.

### 4.2 Perspective

In conclusion, WGS is an effective ”all-in-one” method for detecting the most common abnormalities in cell lines, and validating on-target edits. We agree with the sentiment expressed in a 2020 paper about aneuploidy detection by Assou *et al*., who wrote: ”Ultimately, we anticipate that NGS will become the primary technique for assessing hPSC quality because sequencing costs continue to decrease.”^1^We believe that SeqVerify will unlock the potential of WGS for hPSC quality control, and more broadly, for verifying the quality of any cell lines.

### 4.3 Limitations of the study

The main limitations of SeqVerify are based on its use of short-read DNA sequencing as input. SeqVerify is not set up to detect balanced translocations, and in the case of different transgenes with shared terminal regions, it cannot tell which transgene is present at each insertion site. Long-read sequencing would be a better option in those cases. Furthermore, epigenetic or transcriptional abnormalities cannot be detected based only on DNA sequence data. Notably, PSCs can have epigenetic abnormalities such as loss of imprinting^28^, or erosion of X-inactivation for female PSC lines^29^. Since these abnormalities do not alter the genomic DNA sequence, they are not currently detectable by short-read WGS technology, so alternative methods should be used to detect them. We are particularly interested in using nanopore sequencing to directly detect DNA methylation for this purpose.

## Supporting information

Supplementary File 1: SeqVerify usage guide

Supplementary File 2: KRAKEN reports

Supplementary File 3: Example output

Supplementary File 4: Plasmid sequences

Supplementary File 5: Table of primers

Supplementary File 6: Gel images

## 5 Acknowledgements

Funding for this project was provided by an NICHD F31 fellowship to Merrick Pierson Smela (F31HD108898-01A1). Portions of this research were conducted on the O2 High Performance Compute Cluster, supported by the Research Computing Group, at Harvard Medical School. We thank Dr. Chun-Ting Wu for assistance with validating *Mycoplasma* detection.

## 6 Author Contributions

Conceptualization, Merrick Pierson Smela (M.P.S); Methodology, M.P.S and Valerio Pepe (V.P); Software, M.P.S and V.P; Validation, M.P.S and V.P; Investigation, M.P.S and V.P; Resources, M.P.S and George M. Church (G.M.C); Writing – Original Draft, M.P.S and V.P; Writing – Review & Editing, M.P.S and V.P; Supervision, G.M.C; Funding Acquisition, M.P.S., G.M.C.

## 9 STAR METHODS

### 9.1 Resource Availability

#### 9.1.1 Lead contact

Further information and requests for resources should be directed to and will be fulfilled by the lead contact, George M. Church.

#### 9.1.2 Materials availability

This study did not generate new unique reagents.

#### 9.1.3 Data and code availability

- All code used in this study is available on Github at: https://github.com/mpiersonsmela/seqverify.
- Raw sequencing reads for hiPSC lines derived from PGP1 are available through the NCBI Sequence Read Archive: PRJNA1019637. In order to respect donor privacy, sequencing data from other hiPSC lines are not publicly available. Sequences of plasmids used for generating knock-ins are provided in Supplementary File 4

### 9.2 Experimental Model Details

#### 9.2.1 iPSC culture

Human iPSCs were cultured in mTeSR Plus medium (Stemcell Technologies) on standard polystyrene culture plates coated with Matrigel (Corning) or Geltrex (Thermo Fisher). Four lines were used: PGP1 (male) and ATCC-BSX0115, ATCC-BSX0116, and F66 (all female). Cells were passaged as small clumps using 0.5 mM EDTA and treated with 10 mM Y-27632 for 24 hr after passage. Cells were routinely tested for *Mycoplasma* using the ATCC Universal Mycoplasma Detection PCR kit. All tested negative, except one sample known to be contaminated with *Mycoplasma* that was used solely as a positive control for the KRAKEN2 analysis.

#### 9.2.2 Generation of knock-in iPSC lines

Generation of knock-in lines was performed as previously described ^30^. Briefly, homology donor plasmids were constructed by Gibson assembly of a bacterial plasmid backbone, 5’ and 3’ homology arms PCR-amplified from genomic DNA, and a fluorescent protein insert. For all knock-ins except *SYCP3*, the insert also contained a PGK-PuroTK selection marker flanked by Rox sites, which was excised upon expression of Dre recombinase. sgRNA oligos targeting the knock-in site were cloned into pX330 (Addgene #42230), which expressed the sgRNA and Cas9. Plasmid sequences are given in Supplementary File 4.

To generate each line, homology donor plasmid and sgRNA/Cas9 plasmid (1 µg each) were co-electroporated into 200,000 hiPSCs using the Lonza 4D nucleofector system with 20 µL of P3 buffer and pulse setting CA-137. The cells were plated in one well of a 6-well plate and, for all knockins except *SYCP3*, selection was begun with puromycin after 48 hours. Subsequently, colonies were picked manually with a P20 pipette, transferred to a 96-well plate, and expanded. If required, a further round of electroporation was performed with pCAGGS-Dre to excise the PuroTK selection marker. During this step, selection was performed with ganciclovir (4 µM).

Preliminary genotyping was performed by PCR (primers sequences and gel images are provided in Supplementary Files 5 and 6), and editing was further confirmed by whole genome sequencing as described below.

#### 9.2.3 DNA extraction and sequencing

Genomic DNA was extracted from hiPSCs using the Qiagen DNeasy Blood and Tissue kit. 1 million hiPSCs were used per sample. Extracted DNA was submitted to Novogene Corporation for library preparation and Illumina whole genome sequencing (150 bp paired-end reads to 10X coverage).

#### 9.2.4 SeqVerify Analysis

SeqVerify analysis was performed for each sample using default settings. Briefly, an augmented reference genome was generated from T2T-CHM13v2.0 containing desired edits as well as transgene sequences for insertion site detection. Reads were aligned to this augmented reference genome using BWA-MEM to generate a BAM file for validation of on-target editing and detection of transgene insertion sites. The BAM file was then passed as input to CNVpytor for CNV analysis. Unaligned reads were passed as input to KRAKEN2 for microbial contaminant detection. For SNV analysis, reads were aligned to the GRCh38 reference genome and variants were called using `bcftools mpileup` and `bcftools call`. Variants were filtered by quality score and annotated using snpEff and snpSift and the ClinVar reference database.

A full description of the SeqVerify pipeline, including instructions on usage, is provided in Supplementary File 1.

## References

1. Assou, S. et al. Recurrent Genetic Abnormalities in Human Pluripotent Stem Cells: Definition and Routine Detection in Culture Supernatant by Targeted Droplet Digital PCR. Stem Cell Reports 14, 1–8 (2020).

2. Taapken, S. M. et al. Karyotypic abnormalities in human induced pluripotent stem cells and embryonic stem cells. Nat Biotechnol 29, 313–314 (2011).

3. Markouli, C. et al. Gain of 20q11.21 in Human Pluripotent Stem Cells Impairs TGF-β-Dependent Neuroectodermal Commitment. Stem Cell Reports 13, 163–176 (2019).

4. Nguyen, H. T. et al. Gain of 20q11.21 in human embryonic stem cells improves cell survival by increased expression of Bcl-xL. MHR: Basic science of reproductive medicine 20, 168–177 (2014).

5. Merkle, F. T. et al. Human pluripotent stem cells recurrently acquire and expand dominant negative P53 mutations. Nature 545, 229–233 (2017).

6. Rouhani, F. J. et al. Substantial somatic genomic variation and selection for BCOR mutations in human induced pluripotent stem cells. Nat Genet 54, 1406–1416 (2022).

7. Thompson, O. et al. Low rates of mutation in clinical grade human pluripotent stem cells under different culture conditions. Nat Commun 11, 1528 (2020).

8. Merkle, F. T. et al. Whole-genome analysis of human embryonic stem cells enables rational line selection based on genetic variation. Cell Stem Cell 29, 472-486.e7 (2022).

9. Olarerin-George, A. O. & Hogenesch, J. B. Assessing the prevalence of mycoplasma contamination in cell culture via a survey of NCBI’s RNA-seq archive. Nucleic Acids Research 43, 2535–2542 (2015).

10. Simkin, D. et al. Homozygous might be hemizygous: CRISPR/Cas9 editing in iPSCs results in detrimental on-target defects that escape standard quality controls. Stem Cell Reports 17, 993–1008 (2022).

11. Veres, A. et al. Low Incidence of Off-Target Mutations in Individual CRISPR-Cas9 and TALEN Targeted Human Stem Cell Clones Detected by Whole-Genome Sequencing. Cell Stem Cell 15, 27–30 (2014).

12. McGrath, E. et al. Targeting specificity of APOBEC-based cytosine base editor in human iPSCs determined by whole genome sequencing. Nat Commun 10, 5353 (2019).

13. Zuo, E. et al. Cytosine base editor generates substantial off-target single-nucleotide variants in mouse embryos. Science 364, 289–292 (2019).

14. Wood, D. E., Lu, J. & Langmead, B. Improved metagenomic analysis with Kraken 2. Genome Biol 20, 257 (2019).

15. Landrum, M. J. et al. ClinVar: improving access to variant interpretations and supporting evidence. Nucleic Acids Res 46, D1062–D1067 (2018).

16. Rhie, A. et al. The complete sequence of a human Y chromosome. Nature 621, 344–354 (2023).

17. Nurk, S. et al. The complete sequence of a human genome. (2022).

18. Li, H. & Durbin, R. Fast and accurate short read alignment with Burrows-Wheeler transform. Bioinformatics 25, 1754–1760 (2009).

19. Thorvaldsdóttir, H., Robinson, J. T. & Mesirov, J. P. Integrative Genomics Viewer (IGV): high-performance genomics data visualization and exploration. Brief Bioinform 14, 178–192 (2013).

20. Suvakov, M., Panda, A., Diesh, C., Holmes, I. & Abyzov, A. CNVpytor: a tool for copy number variation detection and analysis from read depth and allele imbalance in whole-genome sequencing. Gigascience 10, giab074 (2021).

21. Peterson, S. E. & Loring, J. F. Genomic Instability in Pluripotent Stem Cells: Implications for Clinical Applications. Journal of Biological Chemistry 289, 4578–4584 (2014).

22. Lu, J., Breitwieser, F. P., Thielen, P. & Salzberg, S. L. Bracken: estimating species abundance in metagenomics data. PeerJ Computer Science 3, e104 (2017).

23. Li, H. A statistical framework for SNP calling, mutation discovery, association mapping and population genetical parameter estimation from sequencing data. Bioinformatics 27, 2987–2993 (2011).

24. Cingolani, P. et al. A program for annotating and predicting the effects of single nucleotide polymorphisms, SnpEff: SNPs in the genome of Drosophila melanogaster strain w ^1118^; iso-2; iso-3. Fly 6, 80–92 (2012).

25. Cingolani, P. et al. Using Drosophila melanogaster as a Model for Genotoxic Chemical Mutational Studies with a New Program, SnpSift. Front. Gene. 3, (2012).

26. Sherman, E. et al. INSPIIRED: A Pipeline for Quantitative Analysis of Sites of New DNA Integration in Cellular Genomes. Molecular Therapy - Methods & Clinical Development 4, 39–49 (2017).

27. Warburton, D. De Novo Balanced Chromosome Rearrangements and Extra Marker Chromosomes identified at Prenatal Diagnosis: Clinical Significance and Distribution of Breakpoints.

28. Bar, S., Schachter, M., Eldar-Geva, T. & Benvenisty, N. Large-Scale Analysis of Loss of Imprinting in Human Pluripotent Stem Cells. Cell Reports 19, 957–968 (2017).

29. Cloutier, M. et al. Preventing erosion of X-chromosome inactivation in human embryonic stem cells. Nat Commun 13, 2516 (2022).

30. Pierson Smela, M. D. et al. Directed differentiation of human iPSCs to functional ovarian granulosa-like cells via transcription factor overexpression. eLife 12, e83291 (2023).

