## Supplementary File 1: SeqVerify usage guide for "SeqVerify: An accessible analysis tool for cell line genomic integrity, contamination, and gene editing outcomes"

### 1. Running SeqVerify

SeqVerify only includes the `seqverify` command, so all calls through the package are done through providing different options. The following command is the minimal SeqVerify call with both exact and inexact (untargeted) insertions:

```
seqverify --reads_1 R1.fastq
--reads_2 R2.fastq --inexact transgenes.fa --exact commands.txt
```

This requires the forward and reverse reads in FASTQ format, a FASTA file containing the sequences that should be reported on in the insertion site detection portion of the pipeline, and a TXT file formatted as a command file to specify what exact insertions were made such that the reference genome can be altered. The command above will not run the KRAKEN2 and variant calling portions of the pipeline, since those options are not enabled by default.

The SeqVerify options, and their default settings, are listed below:

#### 1.2 General Options

- `--output` sets the name of the sample, affecting most output filenames and the folder name. It is set to `output` by default, setting the output folder to be named `seqverify_output`.
- `--reads_1` and `--reads_2` set the paired-read FASTQ (or gzipped FASTQ) source files. Also accepts paths to the files if they're not in the working directory (e.g. for use in research clusters).
- `--threads` and `--max_mem` regulate performance: the former sets how many threads should be used by the pipeline, the latter puts a cap on memory in the pipeline's most memory-intensive process, Burrows-Wheeler alignment, as well as the Java-based subprocesses that the pipeline uses.
- `--start` allows the user to start and stop at any point in the pipeline; this can be done for core efficiency (if on a job scheduler, run all the single-threaded portions on one job, and all the multi-threaded portions on another), as well as the event that parts of the pipeline have already been run but more analysis is desired, or in the event of an error such as the machine turning off, allows for the pipeline to pick back up where it left off. The valid options for `--start` are:
    - o "`beginning`", the default option, which runs the entire pipeline from the start.
    - o "`align`", which skips the creation of the augmented genome, starting at the alignment process (useful for cohorts of samples where the user is looking for the same markers/transgenes on the same reference genome).

- o "`markers`", which skips to the creation of the insertion site readout (useful if the readout was skipped in a previous run and the user now desires it).
- o "`cnv`", which skips to the CNV analysis portion of the pipeline (useful if the user is not interested in the insertion site detection or skipped it on a previous run).
- o "`plots`", which skips to the CNV plot generation (useful if the user wants to refresh their plots without re-running the CNV analysis itself).
- o "`kraken`", which skips to the KRAKEN contamination detection portion of the pipeline (useful if the user is only interested in contamination or if they skipped it on a previous run).
- o "`variant`", which only runs the SNV analysis portion of the pipeline (useful as a separate option due to its resource-intensiveness).

- `--keep_temp` can be set to keep the temp folder if a user wants to keep the temporary files (including the intermediate SAM files produced during alignment, the coverage map used to compute the CNV analysis, and the FASTQ files for all unaligned reads, among other files.) It is off by default, as these files can take up >100 GB per sample.
- `--download_defaults` downloads the default genomes and databases to the working directory. These are T2T-CHM13v2.0 for use in `--genome`, GHRCh38/hg38 for use in SNV analysis, and the 8GB PlusPFP KRAKEN2 database for use with `--kraken`. Kills the program after downloading these. If `--genome` or `--kraken` get left blank, SeqVerify will automatically attempt to retrieve these to use as a default option.

### 1.3 Insertion Site Options

- `--genome` takes in the file name of the reference genome to be used for everything except (usually) SNV analysis. If left blank as above, it uses T2T-CHM13v2.0, the default genome downloaded by the pipeline.
- `--inexact` sets the names (or paths if not in the working directory) of the FASTA files containing the sequences of the markers to detect the insertions sites of (transgenes, plasmids, etc.). Accepts more than one argument, space-separated, if necessary.
- `--exact` is the name or path to a valid command file for insertion of markers where the insertion site is known. Further details on the construction of a valid command file are given below. Only accepts one command file (but a command file can have multiple commands, so this will not restrict analysis).

- --granularity and --min_matches set the insertion site detection parameters: the former regulates how wide the window of a single insertion site is (default: 500), and the latter sets how many matches must be present at a single site for the site to appear in the readout (default: 1, but a higher number may reduce false positive alignments due to repetitive DNA or other factors).

#### 1.4 CNV analysis options

- --manual_plots can be set to use the alternative matplotlib system for plotting transgene CNV

- --bin_size can be used to set the bins for the Manhattan plot (default: 100000).

#### 1.5 KRAKEN options

- --kraken can be set to enable KRAKEN2/BRACKEN analysis, as long as the --database option is also enabled and is followed by a path to a valid KRAKEN2 database (if left blank, SeqVerify will use the default database PlusPF-8GB).

#### 1.6 Variant calling options

- --variant_calling can be set to enable SNV analysis on the sample. It takes two additional arguments: the genome to be used to re-align the reads for SNV analysis, as well as the annotation database to use (if left blank, both of these have default options the pipeline can fall back upon). --variant_intensity sets the lowest severity to be reported in the final readout, out of *MODIFIER, LOW, MODERATE, HIGH*.

- --min_quality sets a minimum quality filter (using the Phred quality scale) for both variant calling and the similarity detection portion of the pipeline. SNPs above this score will be counted and saved, SNPs below it will be ignored. Set to 100 by default.

- --similarity is a three-argument option: it takes in two VCF files and a minimum severity (from the same set as the --variant_calling), and returns the Jaccard similarity of the two files for simpler stem cell line identification, filtering for all SNPs above or at the given severity. It can also be paired with the --min_quality option to additionally filter based on quality.

### 2. Command Files

SeqVerify is set up to take exact gene edit sites as inputs, as well as inexact (usually transgene) edit sites, in making the insertion site readout. While inexact edits can just be specified by providing the FASTA file of the transgene that the user wants to check for (which will be appended to the genome as an extra nucleotide sequence), to place exact reads SeqVerify requires some additional information, such as the location of the edit, and whether the edit is a deletion, insertion, or replacement.

This is done through a "command file", a specially-formatted text file that the `--exact` flag takes as its argument. One command is uniquely specified by the name of the chromosome (or other sequence) where the edit is taking place, the start and end coordinates to be deleted (unless the command is a pure insertion), and the sequence to be inserted (unless the edit is a pure deletion, in which case no sequence is required), and every line corresponds to a separate command. Commands are thus of the form "`CHR:START-END SEQUENCE`", where the whitespace in between `CHR:START-END` and `SEQUENCE` is a tab character.

SeqVerify will delete the bases from START to END exclusive of both (i.e. deleting bases START+1 to END-1). For example, the command `chr2:0-10` will delete the first 10 bases in chr2 and not replace them with anything. If a sequence is specified, SeqVerify will insert it after deleting the bases from base START+1 onwards: for example, `chr5:10-20 GCT` will delete the 11th to 19th bases and place GCT in the 11th, 12th, and 13th slots. A pure insertion with no deletion can be specified by using the same coordinate for both start and end: `chr1:1-1 AGCT` will not delete anything, and insert AGCT after the first base (i.e. positions 2–5).

Should there be multiple commands acting on the same chromosome that may influence one another (such as a command deleting 3 bases but inserting 5, which will shift all other commands after the site of the insertion by two bases), SeqVerify will automatically handle change in the base coordinates such that the user does not need to work out the effect that a command will have on other commands themselves.

#### 3. Interpreting Output

SeqVerify will output a single folder, `seqverify_output` where *output* is the value of the required `--output` argument, the name of the sample. This will contain at most four subdirectories depending on which portions of the pipeline are performed:

- `insertion`, which contains all files related to the insertion site detection:
    - `seqverify_output_markers.bam` and its corresponding index, a BAM file containing the reads aligned to the genome with the addition of the transgene sequences.
    - `seqverify_readout.txt`, the aforementioned readout for insertion site detection.
    - `seqverify_output_collated.fa`, the reference genome augmented with all inexact transgenes as separate chromosomes
- `copy_number`, containing all of the Copy Number Variation files:

  - output.pytor, the CNVPytor binary file, which can be used to generate further plots or further process the CNV data if necessary.
  - output.global.0000.png, the Manhattan plot of the copy number across the genome provided.
  - calls.bin_size.tsv, all of the CNV calls used by CNVPytor
  - Transgene copy number histograms, produced through either IGV or matplotlib, named fig marker.png where *marker* is the name of the transgene as given in the FASTA file.
- kraken, containing the files related to the contamination analysis:
  - classified_seqs_output.kreport, a human-readable report of the microbial sequences detected by KRAKEN2.
  - classified_output.kraken, the KRAKEN2 binary output files used to generate the report.
  - classified_seqs_output_1.fq, classified_seqs_output_2.fq, the FASTQ files containing the sequnences classified by KRAKEN.
  - classified_seqs_output.bracken, the BRACKEN statistical analysis output for the KRAKEN report.
- variant_calling, containing the files related to SNP calling:
  - seqverify_output.ann.vcf, the database-annotated (ClinVar by default) VCF file output of the SNPs found in the reads provided.
  - seqverify_output_variants.tsv, a human-readable file containing information about all mutations above a certain severity and quality threshold.

An example of SeqVerify output (excluding BAM and VCF files due to size limitations) is provided as Supplementary File 3.
