## Supplementary figures and images for "SeqVerify: An accessible analysis tool for cell line genomic integrity, contamination, and gene editing outcomes"

### 2022-08-29_PGP1_DDX4_wt_ins.png

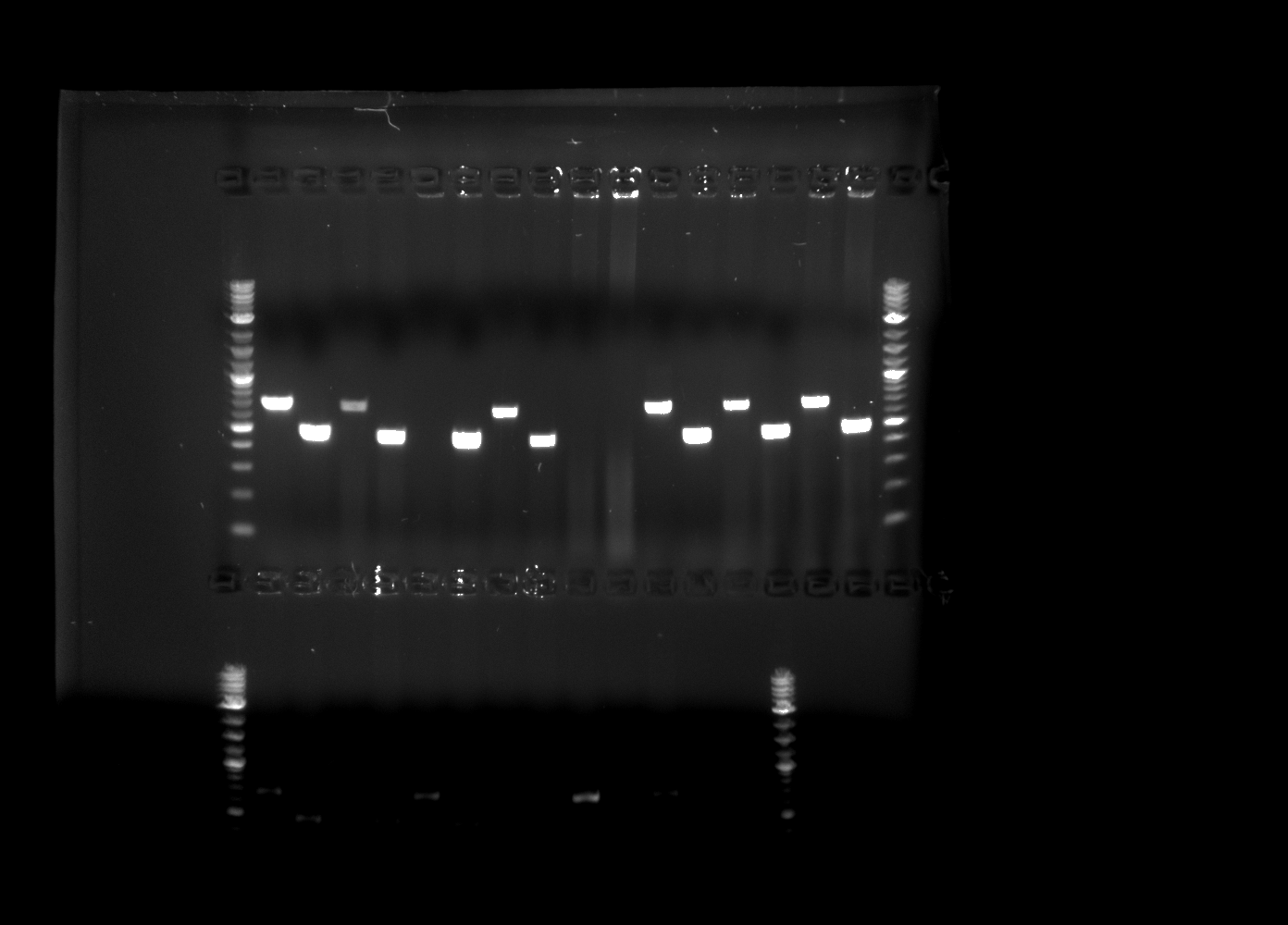

### 2022-08-29_PGP1_DDX4_wt_ins_annotated.png

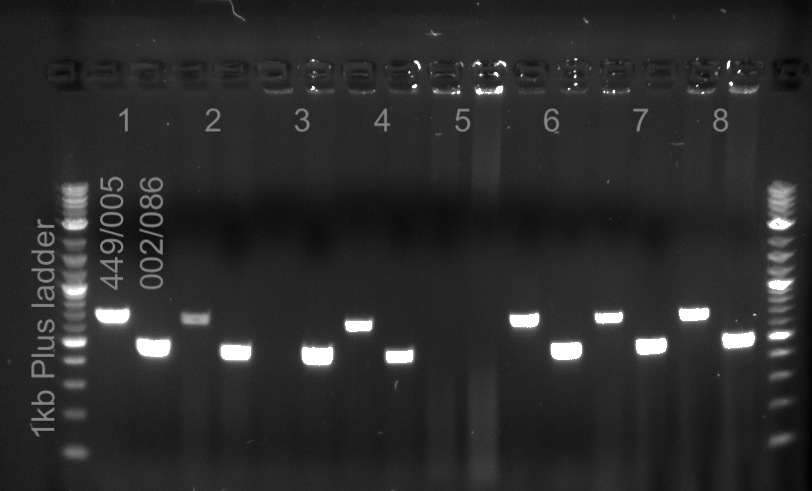

### 2022-11-07_PGP1_3.2_REC8_Dre_021-023+178-024.png

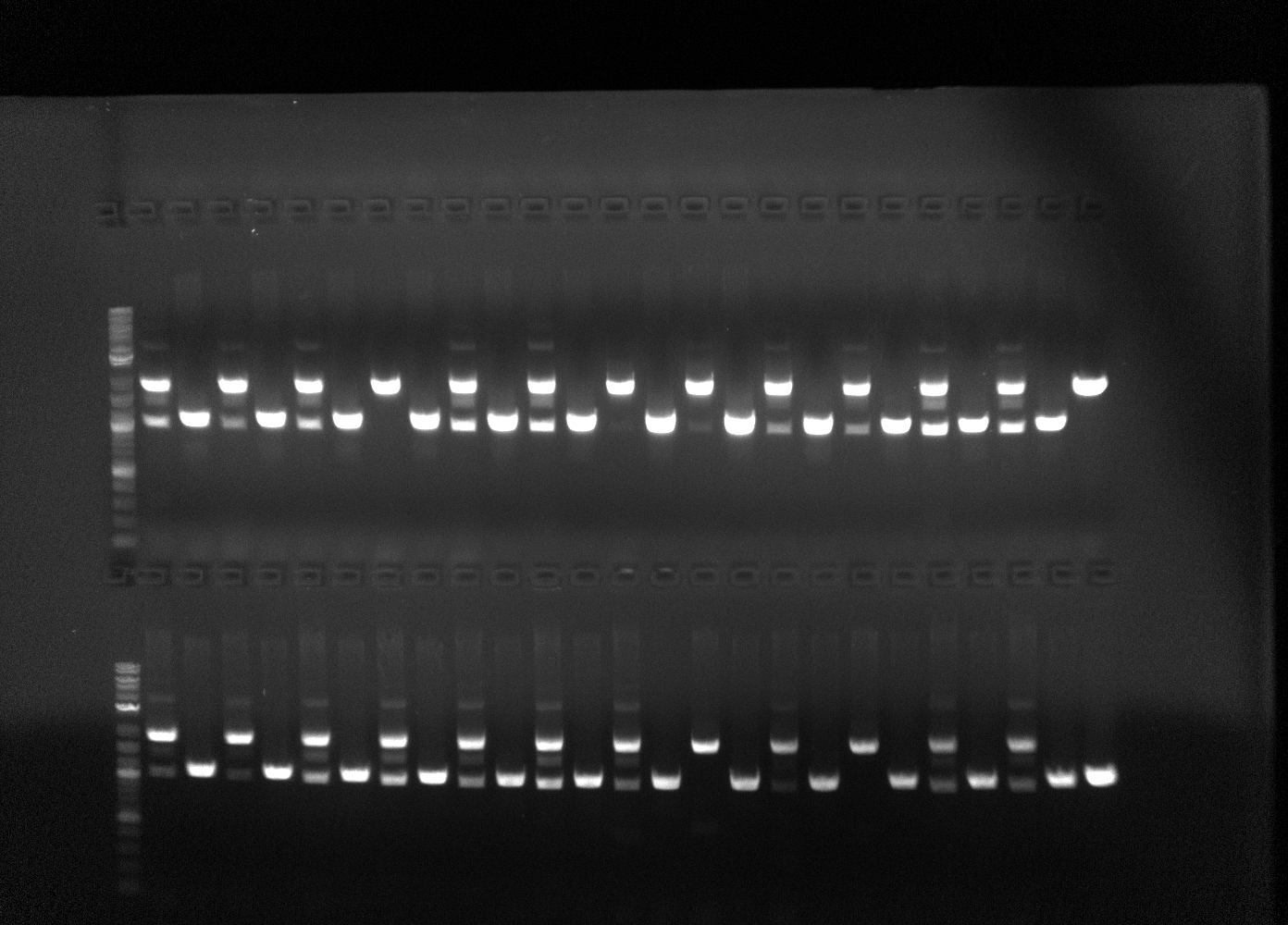

### 2022-11-07_PGP1_3.2_REC8_Dre_021-023+178-024_annotated.png

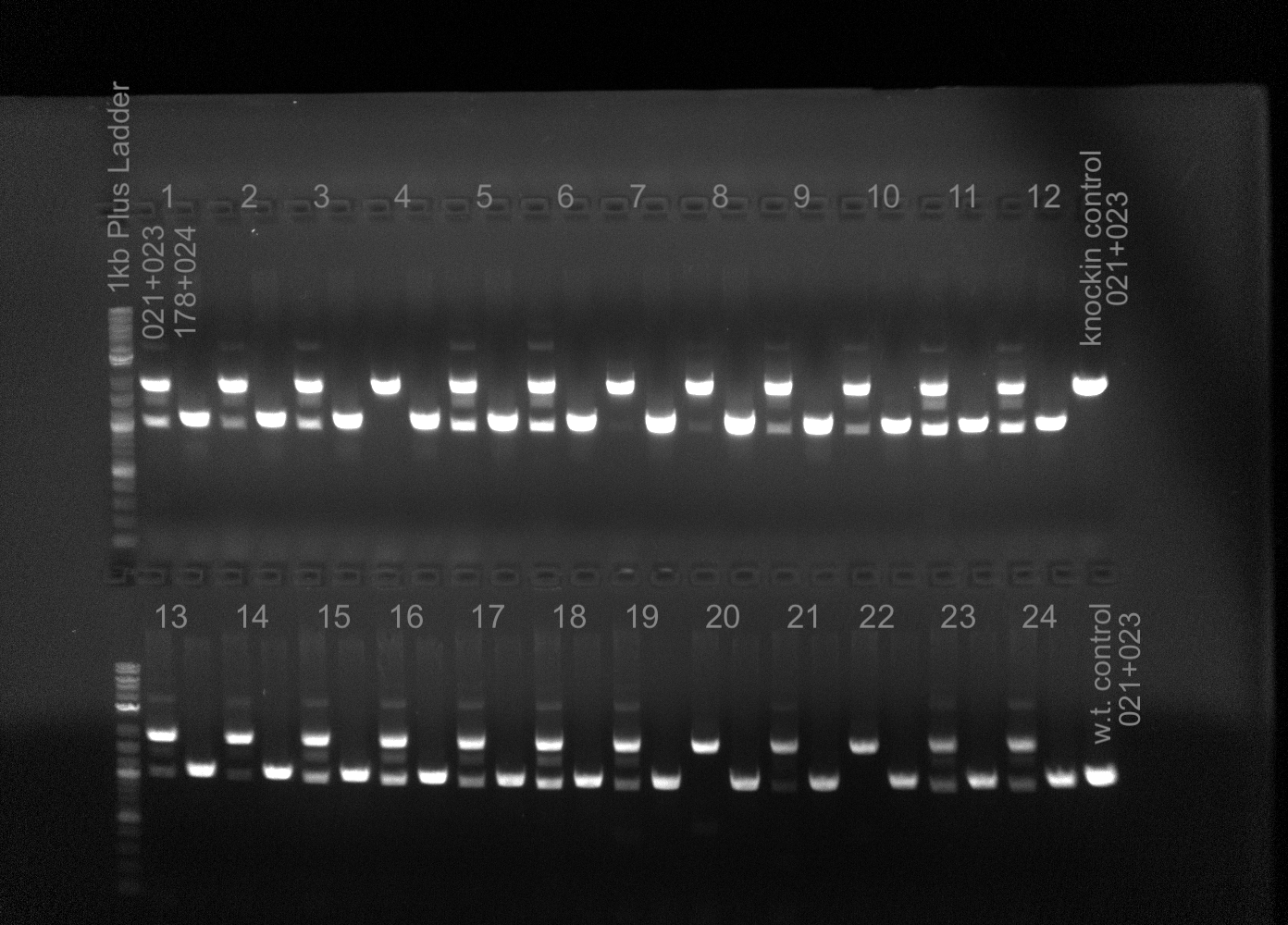

### 2022-12-01_10h30m21s_SYCP3.png

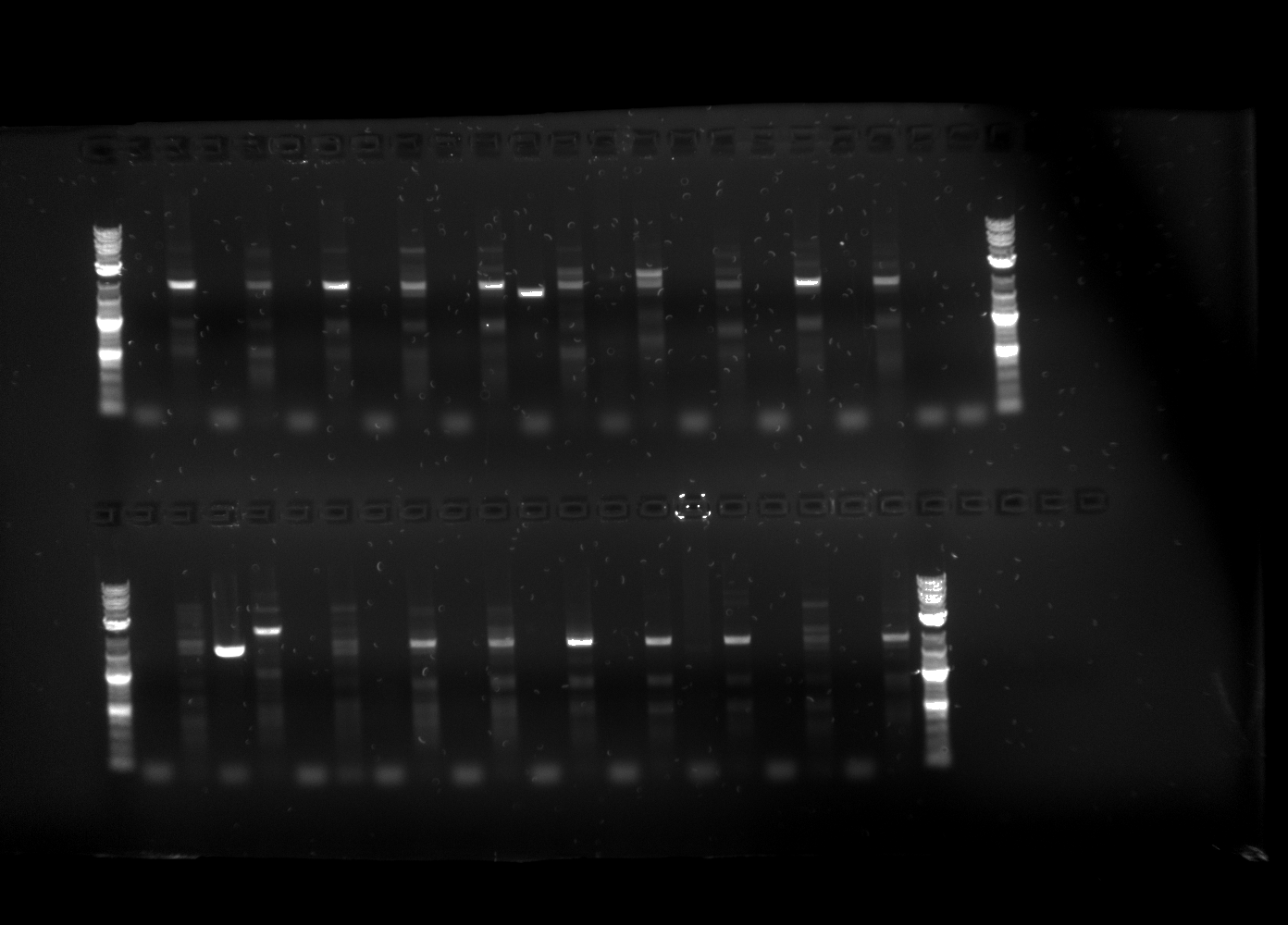

### 2022-12-01_10h30m21s_SYCP3_annotated.png

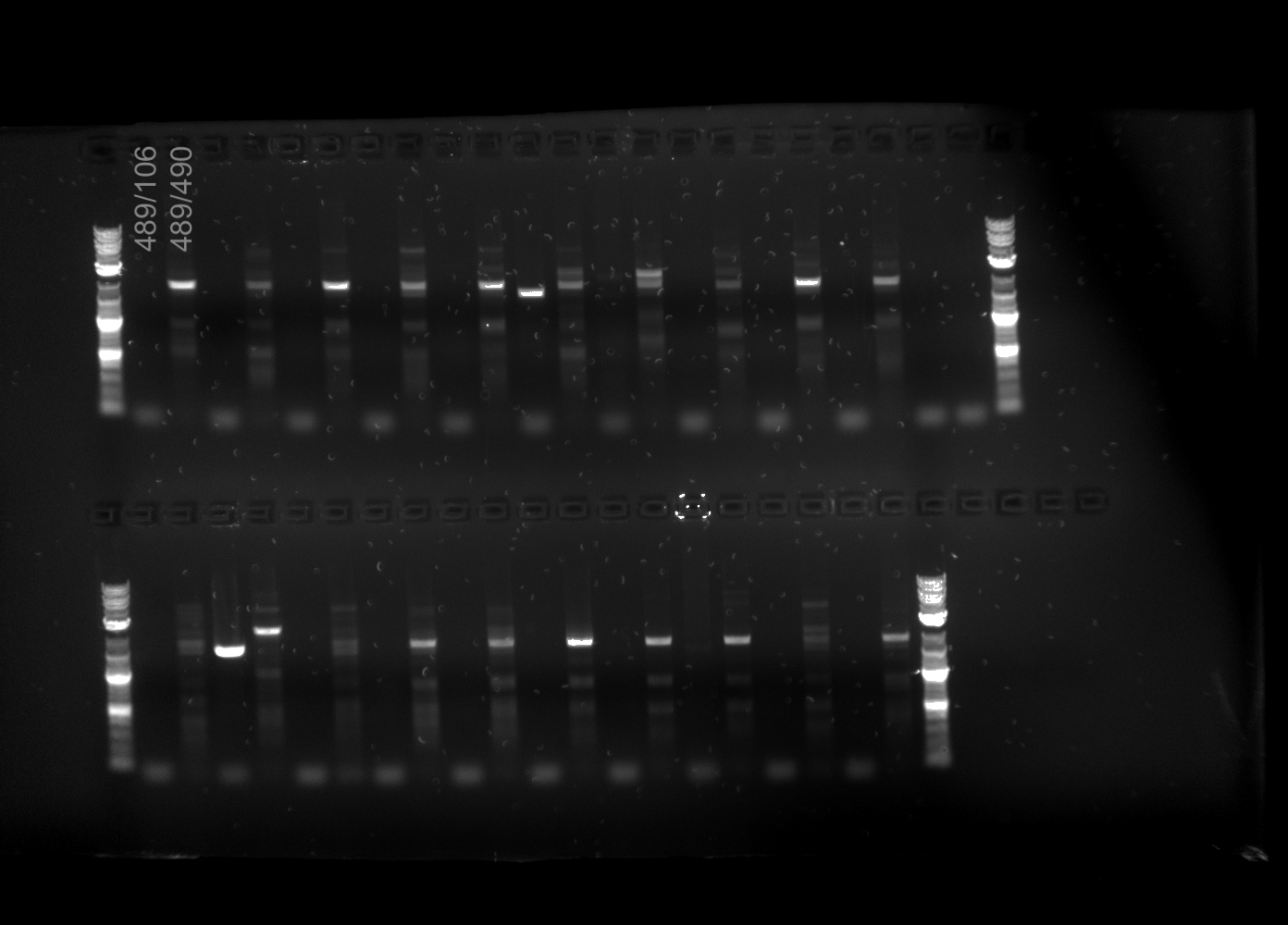

### 2022-12-07_NPM2_NANOS3.png

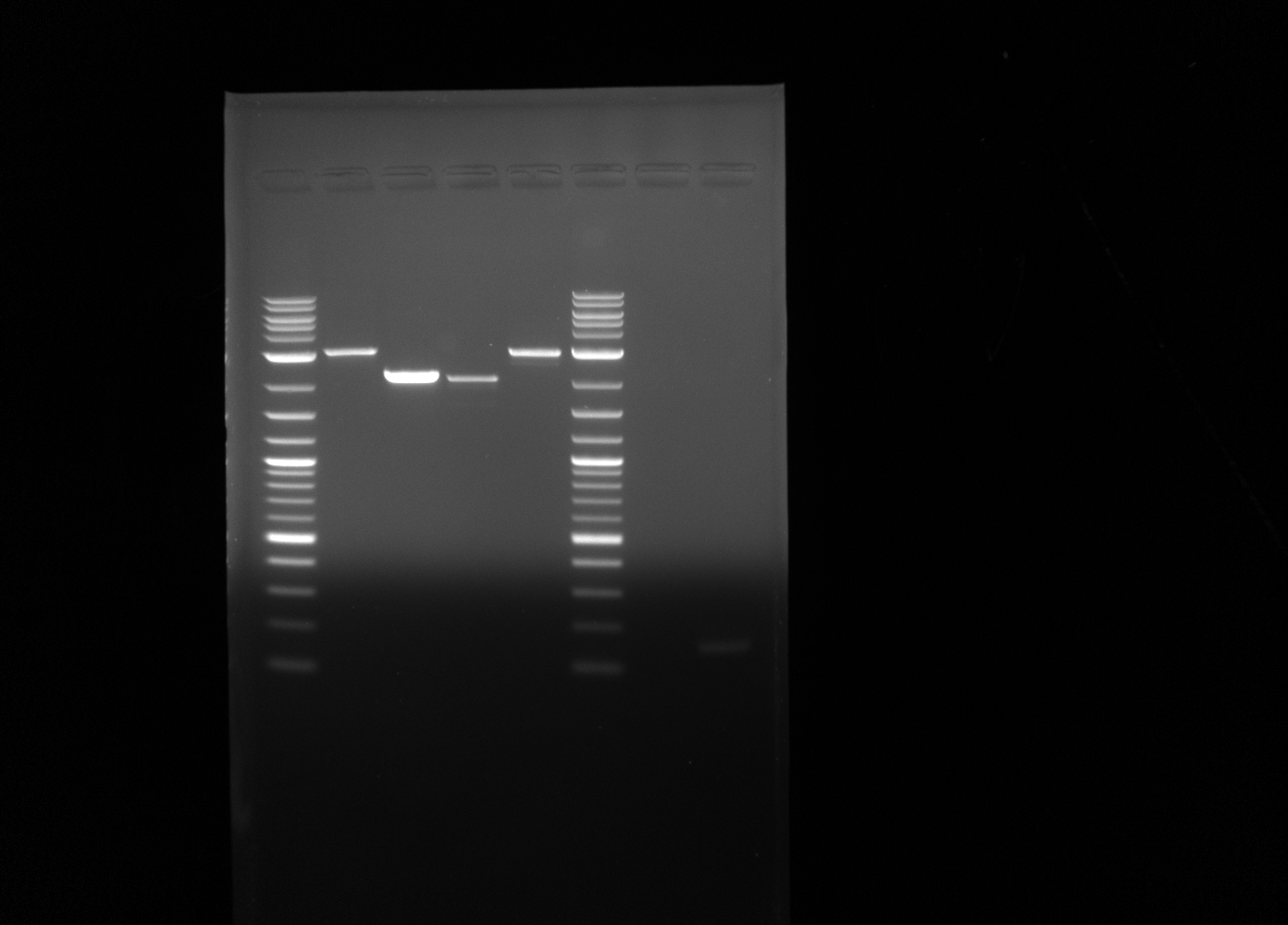

### 2022-12-16_genotyping_fig.png

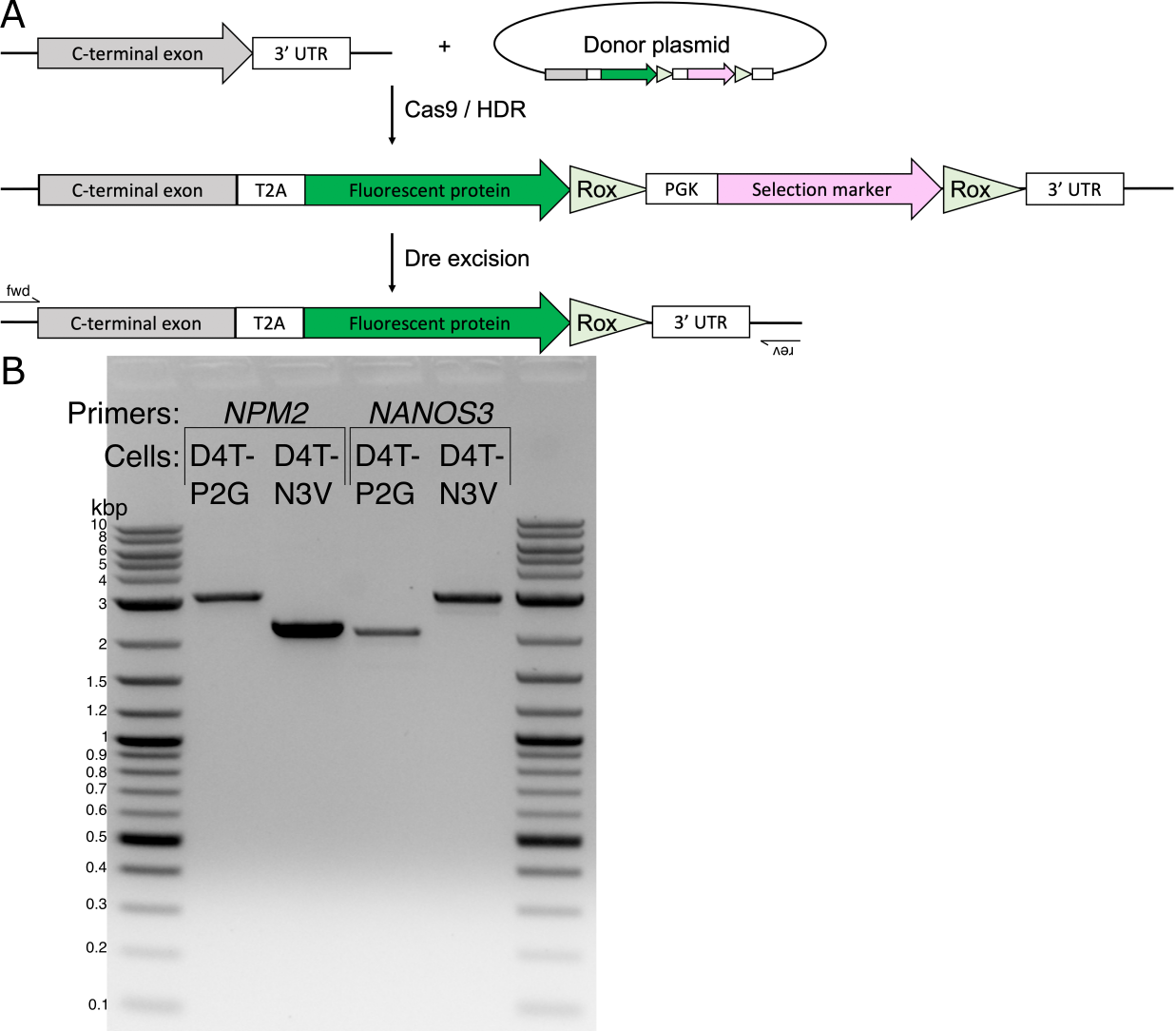

### D_pipetest_2023-09-08.global.0000.png

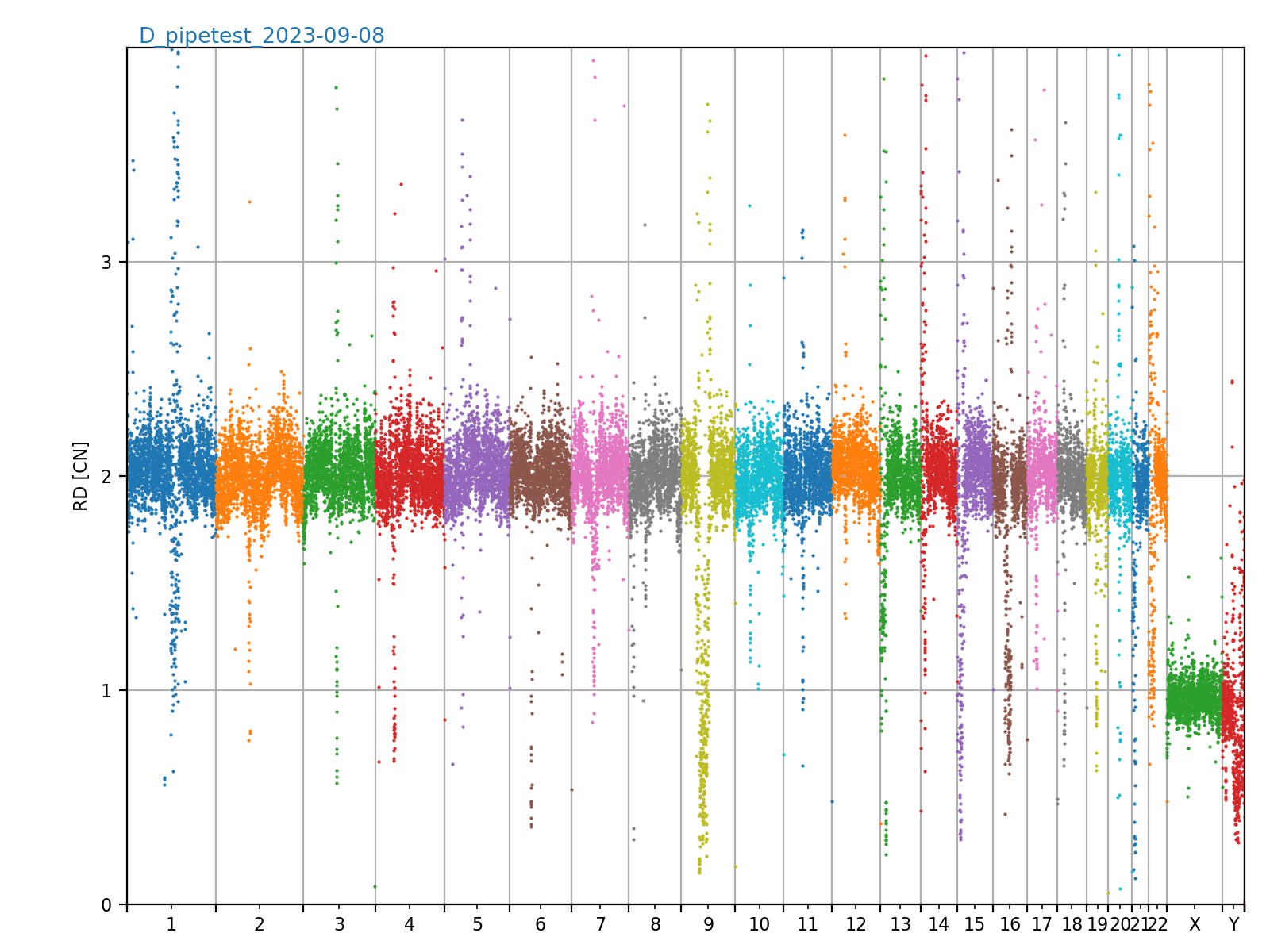

### fig_chr5.png

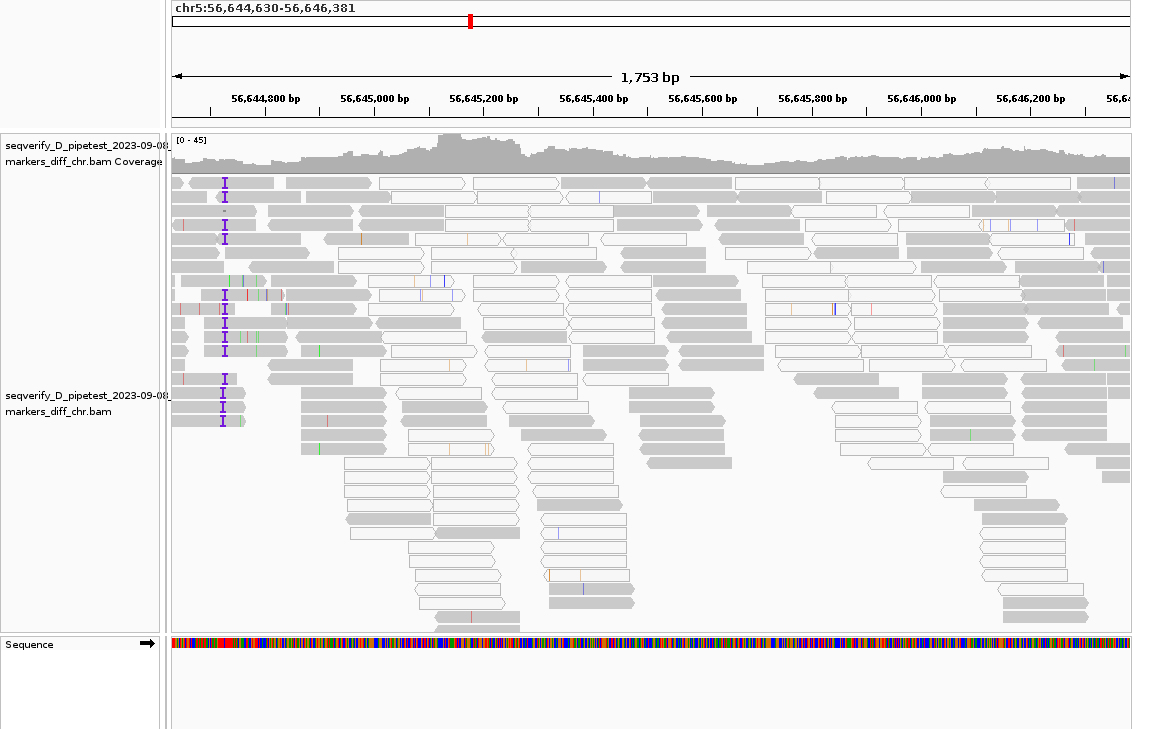

### fig_chr14.png

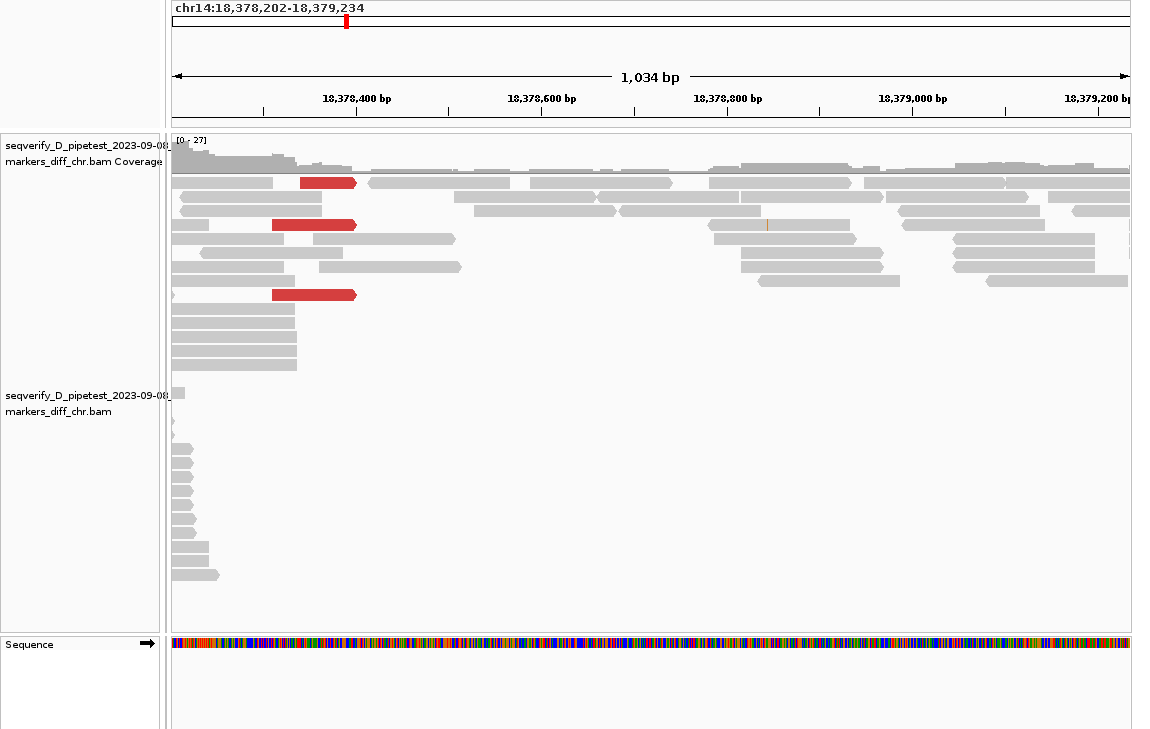

### fig_pCAGGSS-Dre.png

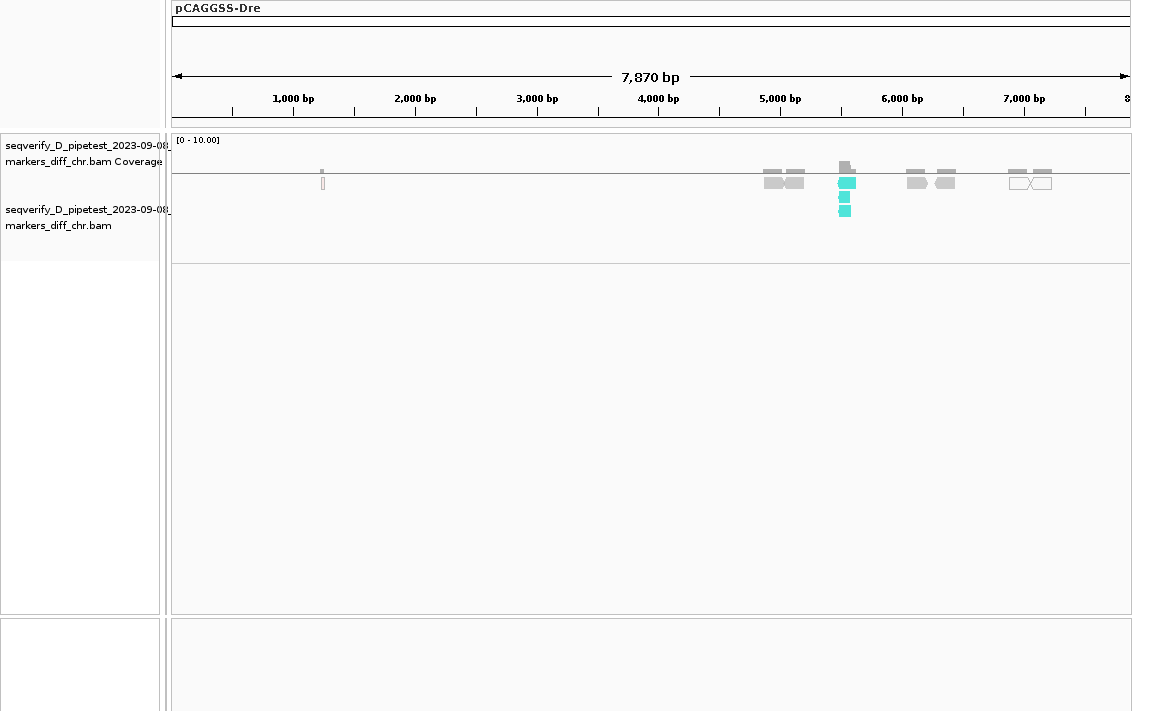

### fig_pHDR2_backbone.png

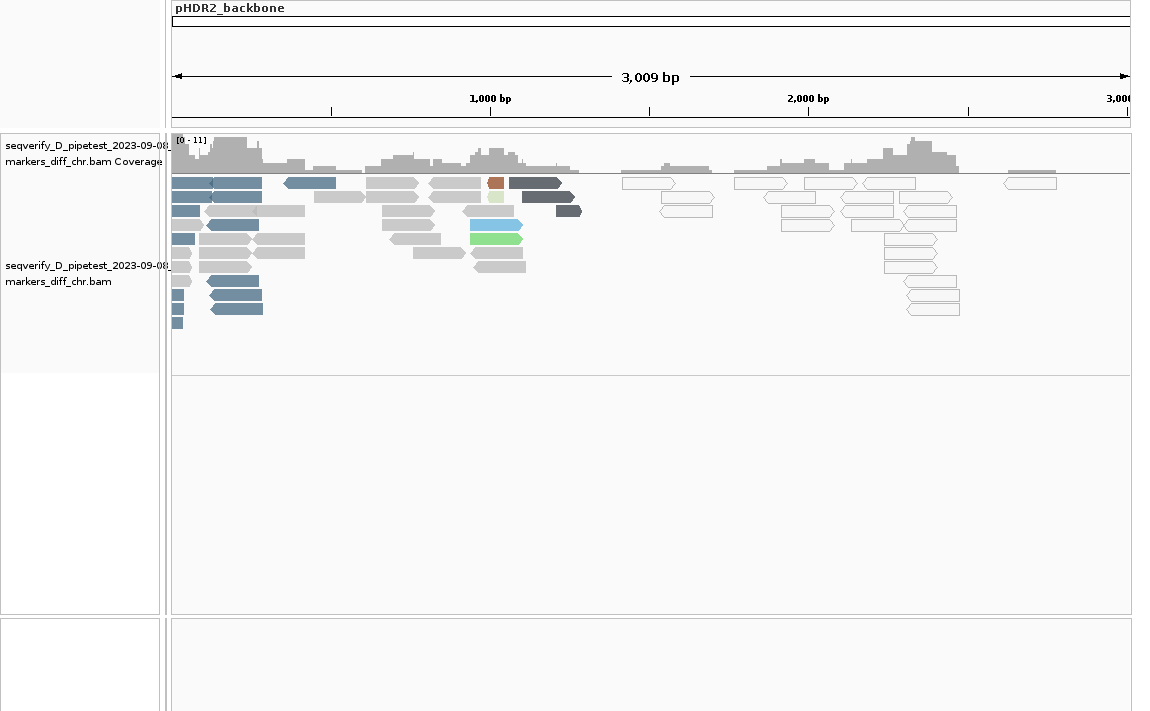

### fig_pX330-U6-Cas9.png

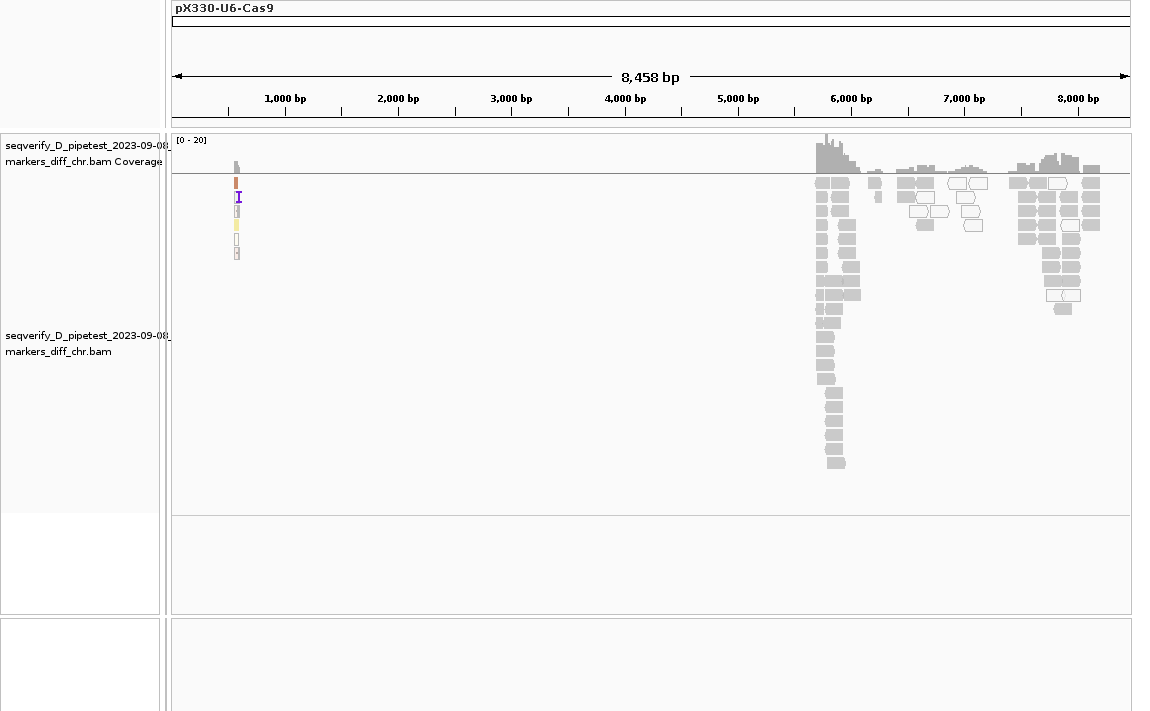

### seqverify_snp_quality.png

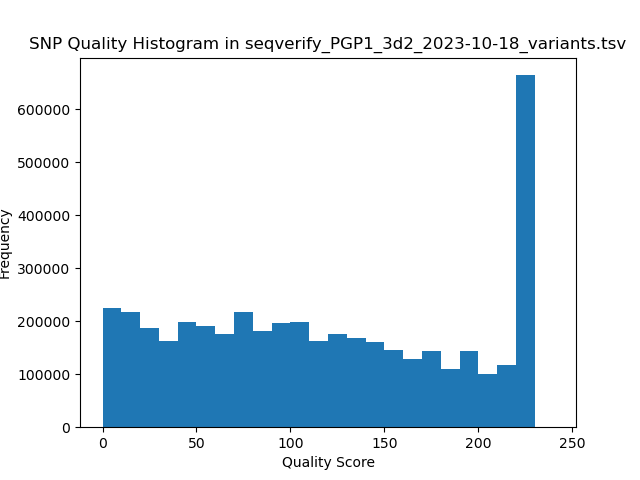
